# Single-step in vitro ribosome reconstitution mediated by two GTPase factors, EngA and ObgE

**DOI:** 10.1101/2024.10.17.618970

**Authors:** Aya Sato, Weng Yu Lai, Yusuke Sakai, Keiko Masuda, Yoshihiro Shimizu

**Affiliations:** Laboratory for Cell-Free Protein Synthesis, RIKEN Center for Biosystems Dynamics research (BDR), Kobe, Hyogo 650-0047, Japan; Graduate School of Frontier Biosciences, The University of Osaka, Osaka 565-0871, Japan; Laboratory of Nucleic Acid Nanotechnology, Institute for Quantitative Biosciences, The University of Tokyo, Tokyo, Japan; Department of Medical Biochemistry and Biophysics, Karolinska Institutet, Stockholm, Sweden

## Abstract

When bacterial ribosomes are assembled in vitro, manipulation of incubation temperature and magnesium ion concentration have been an essential procedure, which is a crucial step for the assembly of active large subunits. The present study tackles with this issue to develop a single-step procedure, which can be performed in a near-physiological condition, where cell-free protein synthesis is active. We found that GTPase factors EngA and ObgE can complement the changes in temperature and magnesium ion concentrations. In the presence of these factors, both the ribosome assembly and translation processes were successfully integrated in the reconstituted cell-free protein synthesis system. Furthermore, we found that these GTPase factors can reassemble the ribosomes to an active state, whose structure was disrupted by EDTA chelation of magnesium ions, indicating that these two factors can reversibly induce the ribosome structure to an intact state. The findings are essential for the bottom-up construction of synthetic cells.

## Introduction

Ribosomes are large intracellular complexes that play a central role in a protein translation system. They consist of two subunits, each composed of one or a few rRNAs and dozens of ribosomal proteins. Maturation processes of these complex assemblies have been studied for decades. For the bacterial ribosomes, the construction of assembly maps of both subunits from *Escherichia coli*, describing the order of ribosomal proteins bind to rRNA, is well known as milestone achievements^1,2^. However, many aspects have yet to be elucidated, including differences between the comparatively slow in vitro reconstitution processes and the more efficient and refined cellular assembly processes, accompanied inclusion of various assembly factors, and effects of a variety of rRNA modifications on these maturation processes. Various in vivo and in vitro studies are still being performed, aiming for a deeper understanding of these processes, by incorporating advanced technologies, including quantitative mass spectrometry (qMS), cryo-electron microscopy (cryo-EM), and single molecule imaging^3-8^.

Recently, with the increasing focus on constructing synthetic cells from the bottom-up viewpoint^9,10^, the importance of the studies on assembling ribosomes from individual parts in vitro has been highlighted. The integrated synthesis, assembly, and translation (iSAT) method developed by Jewett *et al*. demonstrated that in the presence of cell extracts, the assembly of both subunits proceeds using transcribed rRNAs and ribosomal proteins purified from native ribosomes, termed total proteins of the 30S subunit (TP30) and total proteins of the 50S subunit (TP50)^11^. Cell extracts-free ribosome assembly methods have been developed by the integration with the reconstituted cell-free protein synthesis system (PURE system)^12^, enabling the in vitro evolution of 16S rRNA^13^. Also, TP-free methods have been constructed by preparing individually purified recombinant ribosomal proteins for both subunits^14-17^. The use of cell-free expressed ribosomal proteins is also addressed^8,16,18-20^, aiming for the development of self-replicating synthetic cells in the future^21^.

These cell-free synthetic biology studies can help us understand the minimally required building blocks for producing functional assemblies. For the small subunit, functionally active subunits have been shown to be assembled by merely mixing synthetic parts, transcribed rRNA and recombinant ribosomal proteins with minimal post-translational modifications^16^. The functional large subunit has also been shown to be assembled with recombinant proteins with minimal post-translational modifications^17^. However, two major issues exist with the large subunit reconstitution method.

The first issue is the requirement of posttranscriptional modifications in 23S rRNA. According to Green and Noller, modified nucleotides in certain regions of rRNA are necessary for the peptidyl transferase activity of the reconstituted large subunit from *E. coli*^22^. For *Bacillus stearothermophilus*, it has been shown that the subunit with the peptidyl transferase activity can be reconstituted even with the transcribed 23S rRNA^23^. The addition of some chemical substances into the assembly mixtures for *E. coli* ribosomes has also been reported to be effective for the peptidyl transferase activity of the reconstituted subunit using transcribed 23S rRNA^24^. All of these experiments have focused on the peptidyl transferase activity of the reconstituted subunit. Whether or not the assembled subunits are competent for the whole translation processes, including initiation, elongation, and termination steps, remains unclear. By contrast, protein synthesis activity of the reconstituted large subunit was observed if the iSAT method is applied^11^. It can be considered that some factors in the cell extract, including rRNA modification enzymes, are involved in the functional assembly of the large subunit.

The second issue is the need for a two-step procedure that involves temperature and ion concentration changes in the reaction solution. Nierhaus and Dohme revealed that two significant steps are required for the successful assembly of the large subunit^25,26^. In the first step, 5S rRNA, 23S rRNA, and ribosomal proteins are incubated at 44 °C to form 41S or 48S intermediates. Then, by raising the magnesium ion concentration from 4 mM to 20 mM and the temperature to 50 °C, active subunits formation is completed by further incubation (**Fig. 1a**). The need to raise the temperature to a non-physiological value and to manipulate the ion concentrations can be a major barrier when attempting the bottom-up construction of synthetic cells. It is necessary to develop the assembly methods under physiological conditions without temperature changes or ion concentration manipulations. It is noteworthy that the second issue also does not apply to the iSAT method. In the method, the ribosome assembly and the translation processes are seamlessly linked in the same reaction solution, without manipulation of the temperature and ion concentrations^11^. It is highly likely that specific factors in the cell extracts complement this issue.

**Figure 1.**
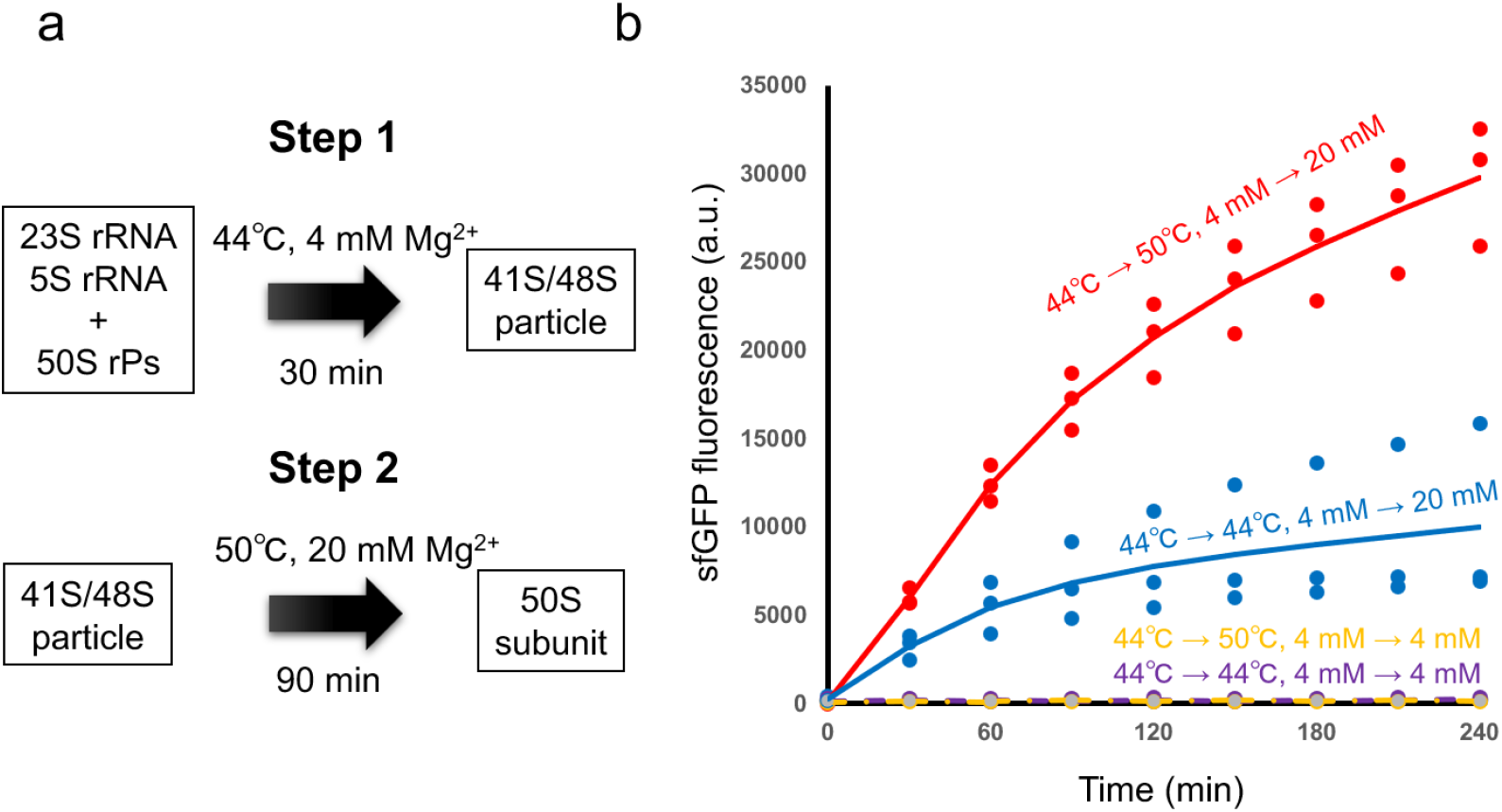
Ribosome assembly with two-step procedure. (a) Schematic of the two-step procedure identified by Nierhaus and Dohme. In the step 1, comparatively low magnesium ion concentration and low incubation temperature are required to assemble the intermediate state of the 50S subunit (41S/48S particle), whereas in the step 2, increasing the incubation temperature and magnesium ion concentration are essential for the assembly of translationally active 50S subunit. (b) Individual effect of temperature and magnesium concentration changes on the activity of assembled ribosomes.

The present study aimed to tackle with the second issue by using ribosome biogenesis factors. It was found that changes in temperatures are effective and changes in magnesium ion concentration are essential for the assembly of translation competent ribosomes. We found that the presence of GTPase factors EngA^27^ and ObgE^28,29^ is necessary to complement these changes. In the presence of these factors, ribosome assembly can proceed under near-physiological conditions, with magnesium ion concentrations below 10 mM, potassium concentrations around 100 mM, and temperatures of 37 °C, where cell-free protein synthesis is active. Both the ribosome assembly and translation processes were successfully linked together in the PURE system, demonstrating that the ribosome assembly proceeds in the same solution conditions as the translation reaction, and the assembled ribosomes are competent for the whole translation processes. Furthermore, it was revealed that unfolded ribosomes, whose structure was disrupted by EDTA chelation of magnesium ions^30^, could be restored to an active state with both the supplementation of magnesium ions and the addition of EngA and ObgE, indicating that these two factors can reversibly induce the ribosome structure to an intact state.

## Results

### Effect of changes in temperature and ion concentration on the ribosome assembly

In our previous study on small subunits, we have found that the assembly proceeds autonomously without the support of any factors^16^, whereas temperature and ion concentration changes are necessary for the assembly of large subunits^25^. To evaluate the impact of these changes on the reconstitution of entire ribosomes, we performed in vitro ribosome assembly experiment using total ribosomal RNAs (5S, 16S, and 23S rRNA) and total proteins of the 70S ribosome (TP70) prepared from the purified native ribosomes^31^. Four patterns of temperature and magnesium ion concentration changes were applied to clarify each effect (**Fig. 1b**). The reconstituted ribosomes were purified from the assembly mixtures and then they were added into the PURE system without the ribosome to test superfolder green fluorescent protein (sfGFP) synthesis. When no magnesium ion concentration changes were applied, sfGFP synthesis by the assembled ribosomes was not detected, and when no temperature changes were applied, sfGFP synthesis decreased by about one-third (**Fig. 1b**). The results showed that the changes in magnesium ions are particularly essential, indicating that the two-step procedure established for the reconstitution of large subunits^25^ (**Fig. 1a**) is also an essential key process for the assembly of entire ribosomes with translation activity, consistent with the previous statement^31^.

### Single-step ribosome assembly using ribosome biogenesis factors

Based on the differences between the iSAT assembly experiments^11^ and those using purified parts, including this study (**Fig. 1b**), we hypothesized that some factors in the cell extracts used in iSAT, particularly ribosome biogenesis factors, may complement the changes in temperature and divalent ion concentration required in the two-step procedure, and facilitate a single-step assembly in the cell-free reaction mixtures (**Fig. 2a**). Furthermore, in previous experiments examining the effects of ribosome biogenesis factors on small subunit reconstitution, we found that the factors Era and RsgA (YjeQ) were effective in promoting subunit assembly, both being GTPase factors^15,16^. Therefore, we focused on GTPase factors, which have been suggested to be involved in large subunit biogenesis. Based on literature surveys, we selected BipA^32^, EngA (Der)^27^, EngB (YihA/YsxC)^33^, HflX^34^, LepA^35^, and ObgE^28,29^, which were prepared as recombinant proteins. Surprisingly, when these six factors were mixed with total ribosomal RNAs (5S, 16S, and 23S rRNA) and TP70, along with an energy regeneration system to maintain GTPase activity, we found that it was possible to assemble the ribosomes with sfGFP synthesis activity at a constant but relatively low magnesium concentration of 8 mM at 37 °C (**Fig. 2b**). The sfGFP synthesis activity significantly dropped at a magnesium concentration of 17 mM in the presence of these six factors. Almost no sfGFP synthesis activity was observed in the absence of six factors, even with 8 mM magnesium, suggesting the selected factors exhibit some activity that compensates for the magnesium ion concentration changes in the two-step procedure. The optimal magnesium concentration for this reaction was 8 mM (**Fig. 3a**), and the optimal monovalent salt concentration for ribosome assembly was several hundred millimolar, consistent with previous assembly experiments^25^ (**Fig. 3b**).

**Figure 2.**
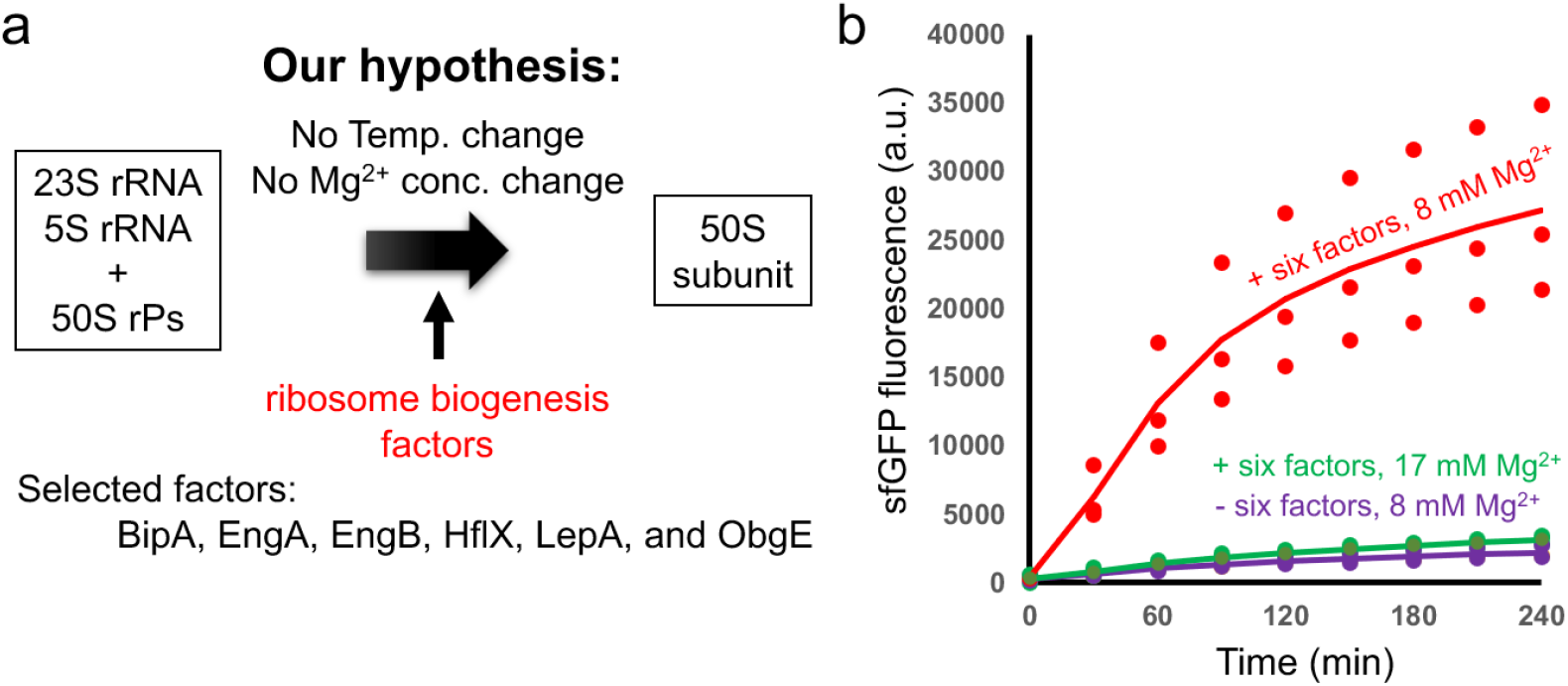
Ribosome assembly with ribosome biogenesis factors. (a) Working hypothesis of this study. We hypothesized that some of the ribosome biogenesis factors take over the role of temperature and magnesium concentration changes in the two-step procedure. We selected six GTPase factors as candidates of these factors. (b) Single-step ribosome assembly by selected ribosome biogenesis factors. Activity of assembled ribosomes in the presence or absence of six factors at 8 or 17 mM magnesium ion concentration at 37 °C. Concentration of potassium glutamate for the ribosome assembly experiment was fixed at 200 mM.

**Figure 3.**
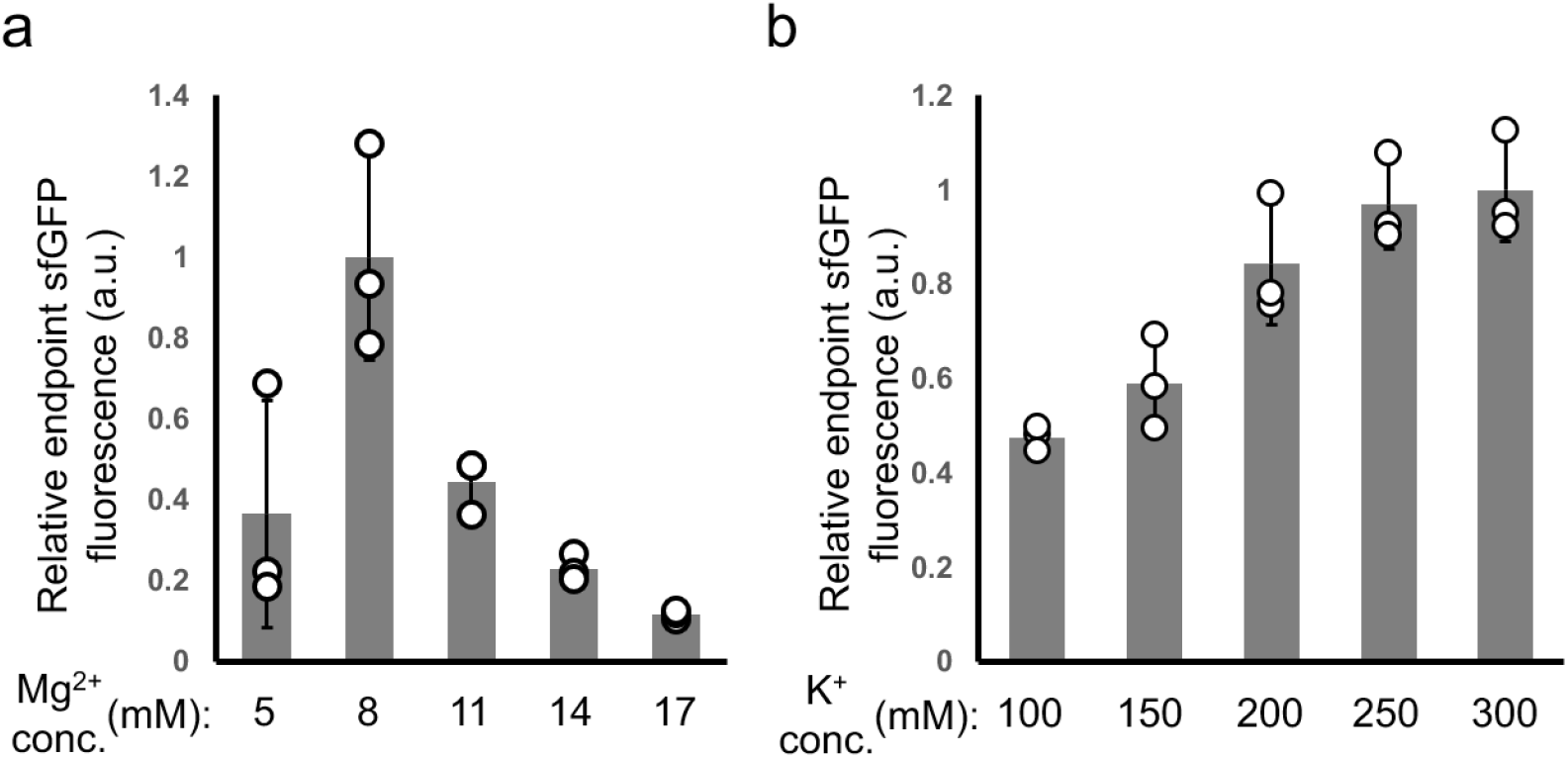
Effect of magnesium and potassium concentrations on the single-step ribosome assembly using six GTPase factors. (a) Effect of magnesium concentration on the assembled ribosome activity. (b) Effect of potassium concentration on the assembled ribosome activity. Relative endpoint sfGFP fluorescence at 4 h timepoint, compared to 8 mM magnesium or 300 mM potassium, respectively, is shown. Potassium concentration was fixed at 200 mM (a) and magnesium concentration was fixed at 8 mM (b).

### Coupling of the assembly and translation reactions

The results with GTPase factors suggested that the factors were able to complement the temperature and ion concentration changes in the two-step procedure, realizing a single-step ribosome assembly. Because the required monovalent and divalent ion concentrations are within the range for the optimum conditions of the translation reactions (**Fig. 3**), it appeared probable to integrate the ribosome assembly and translation processes in a single reaction mixture. To this end, we attempted to construct a coupled reaction, by adding 3 rRNAs, total proteins of the 70S ribosome (TP70), and selected GTPases into the ribosome-omitted PURE system (**Fig. 4a**). The result showed that after a lag period that likely corresponds to the time required for the ribosome assembly, increase of sfGFP fluorescence was observed (**Fig. 4b**). An optimal magnesium concentration was 8 mM (**Fig. 4b, c**), consistent with the large subunit assembly experiment (**Fig. 3a**). The synthesis of sfGFP was unaffected by a monovalent salt concentration above 100 mM (**Fig. 4d**). While high salt concentrations were effective for large subunit assembly (**Fig. 3b**), they may have negatively impacted translation reactions. An empirically translation-competent salt concentration of 100-250 mM^36-38^ was found to be optimal. The results demonstrated that under near-physiological conditions suitable for translation reactions, a single-step ribosome assembly has proceeded in the solution, where the assembled ribosomes are competent for the whole translation processes, resulting in sfGFP synthesis.

**Figure 4.**
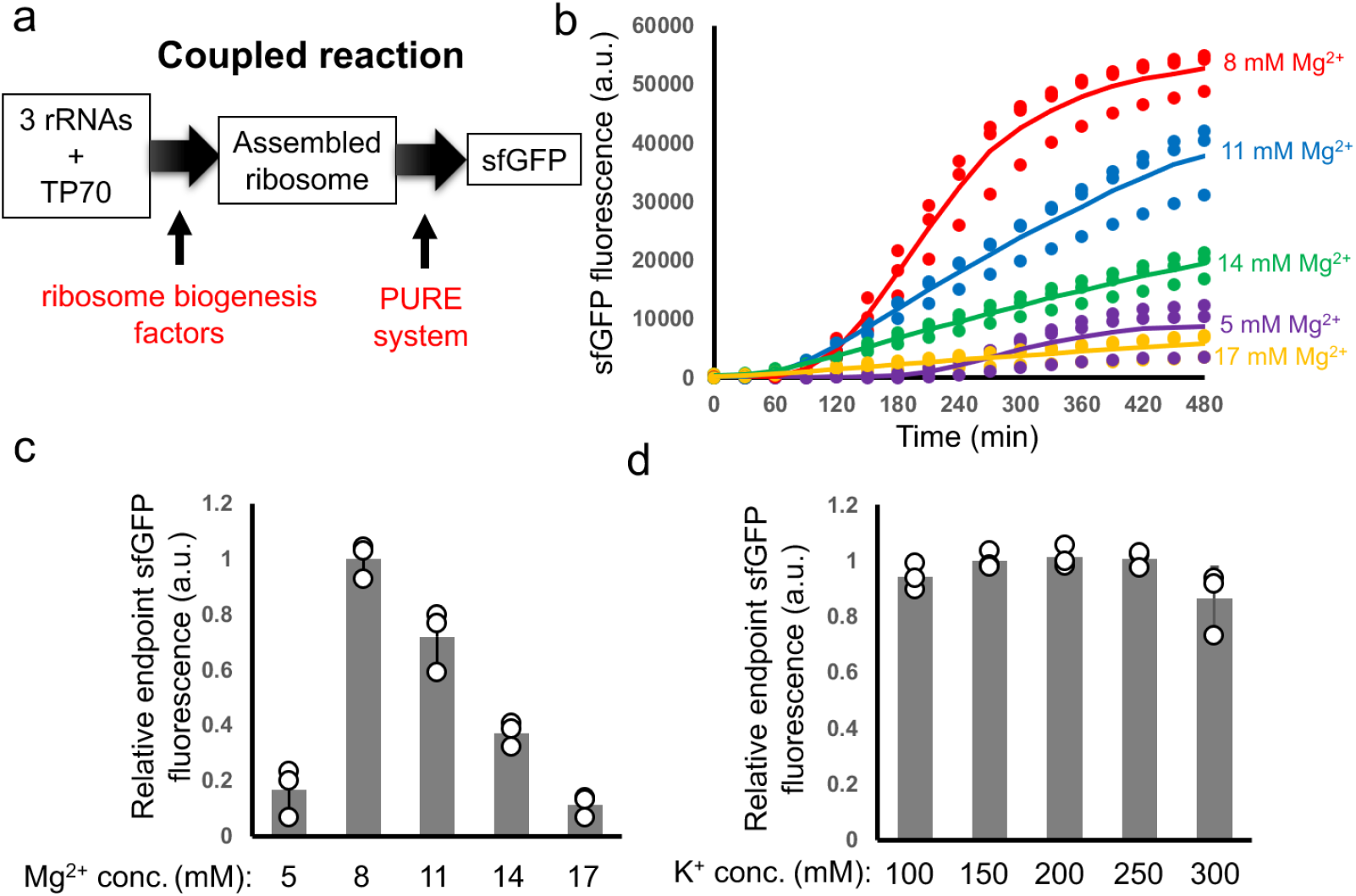
Coupling of the assembly and translation reactions. (a) Schematic of the coupled reaction. Three rRNAs (5S, 16S, and 23S) and TP70 are added into the ribosome-free PURE system. Ribosomes are assembled in the PURE system with a help of ribosome biogenesis factors and then, assembled ribosomes exhibit the translation activity, which can be detected as sfGFP fluorescence. (b) Time course of sfGFP fluorescence in the coupled reaction system. Effect of magnesium concentration is shown. (c) Effect of magnesium concentration on the assembled ribosome activity. (d) Effect of potassium concentration on the assembled ribosome activity. Relative endpoint sfGFP fluorescence at 8 h timepoint, compared to 8 mM magnesium or 150 mM potassium, respectively, is shown. Potassium concentration was fixed at 150 mM (b, c) and magnesium concentration was fixed at 8 mM (d).

### EngA and ObgE are responsible for the single-step ribosome assembly process

We examined which of the selected six factors plays a role in facilitating the single-step ribosome assembly. Since more than one factor could be involved in the process, we addressed the activity of ribosomes assembled using GTPase mixtures with each factor removed one by one. When tested in the coupled reaction system, sfGFP synthesis activity significantly decreased in the absence of ObgE and was not detected in the absence of EngA (**Fig. 5a**). Similar results were obtained in an uncoupled system (**Fig. 5b**), suggesting that EngA and ObgE are the responsible factors which may complement the changes in temperature and ion concentration in the two-step protocol for large subunit assembly. Furthermore, when only ObgE was added to the coupled system, sfGFP synthesis was not detected, similar to the case without any factors, and when only EngA was added, comparatively slow sfGFP synthesis was observed (**Fig. 5c**). When both factors were added, sfGFP synthesis was highly active, indicating that ribosome assembly was achieved with the presence of EngA and ObgE.

**Figure 5.**
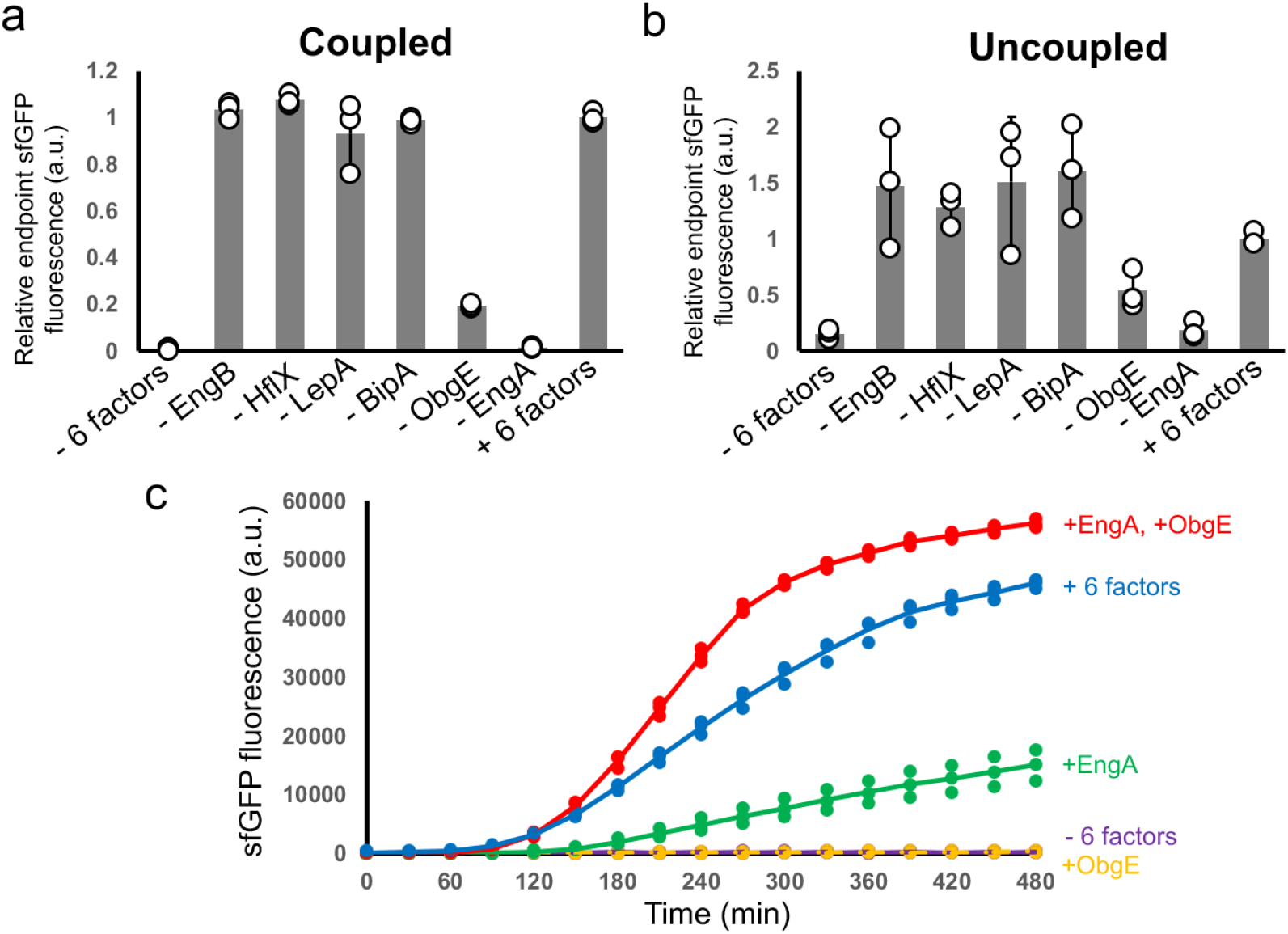
Identification of the responsible factors. (a) Ribosome activities assembled in the absence of each factor in the coupled system. (b) Ribosome activities assembled in the absence of each factor in the uncoupled system. (c) Ribosome activities assembled with only EngA and ObgE. When coupled system is applied, magnesium concentration was fixed at 8 mM and potassium concentration was fixed at 150 mM (a, c). When uncoupled system was used, magnesium concentration was fixed at 8 mM and potassium concentration was fixed at 250 mM for the ribosome assembly reaction (b). Relative endpoint sfGFP fluorescence at 8 h timepoint, compared to that using all six factors (a, b) or time course of sfGFP fluorescence (c) are shown.

Characteristics of the ribosomes assembled using the two factors were analyzed. When comparing the rate of increase in fluorescence of synthesized sfGFP with those of native ribosomes, it was approximately 0.6-fold implying 60% of the input rRNA was assembled into functional ribosomes (**Fig. 6**). SDS-PAGE analysis showed products of the expected size (**Fig. 7a**), and MS analysis allowed assignment of nearly all fragments (**Fig. 7b** and **Supplementary Data 2**), with no apparent issues in fidelity. These results suggest that, although the reconstitution efficiency is around 60%, the ribosomes are reconstituted in a nearly complete form comparable to native ribosomes. We note that with the exception of one peptide fragment detected at an intensity of approximately 1/1000 from +1 frameshift, the MS analysis did not detect any peptide fragments derived from frameshifting or stop-codon readthrough (**Supplementary Data 2)**, suggesting that the reconstituted ribosomes maintain a high level of accuracy in the translation machinery. However, to examine various aspects of translation, such as initiation from internal start codons other than the start codon associated with the SD sequence, or termination at internal stop codons, it will be necessary to select reporters suited to each specific purpose. These issues should therefore be investigated in future studies. In addition to sfGFP, we also targeted dihydrofolate reductase (DHFR) synthesis and confirmed that this protein was produced in an active form (**Fig. 7c**). We note that these results do not indicate that the two factors are necessary and sufficient for ribosome reconstitution. The TP70 used in this study was derived from native ribosomes, and the possibility remains that additional factors are involved. MS analysis of TP70 identified various biogenesis factors, including RsfS, which has been previously reported to function with ObgE in cryo-EM studies^28^ (**Supplementary Data 3**). Further studies will be required to elucidate the detailed mechanisms, including the roles of these additional factors.

**Figure 6.**
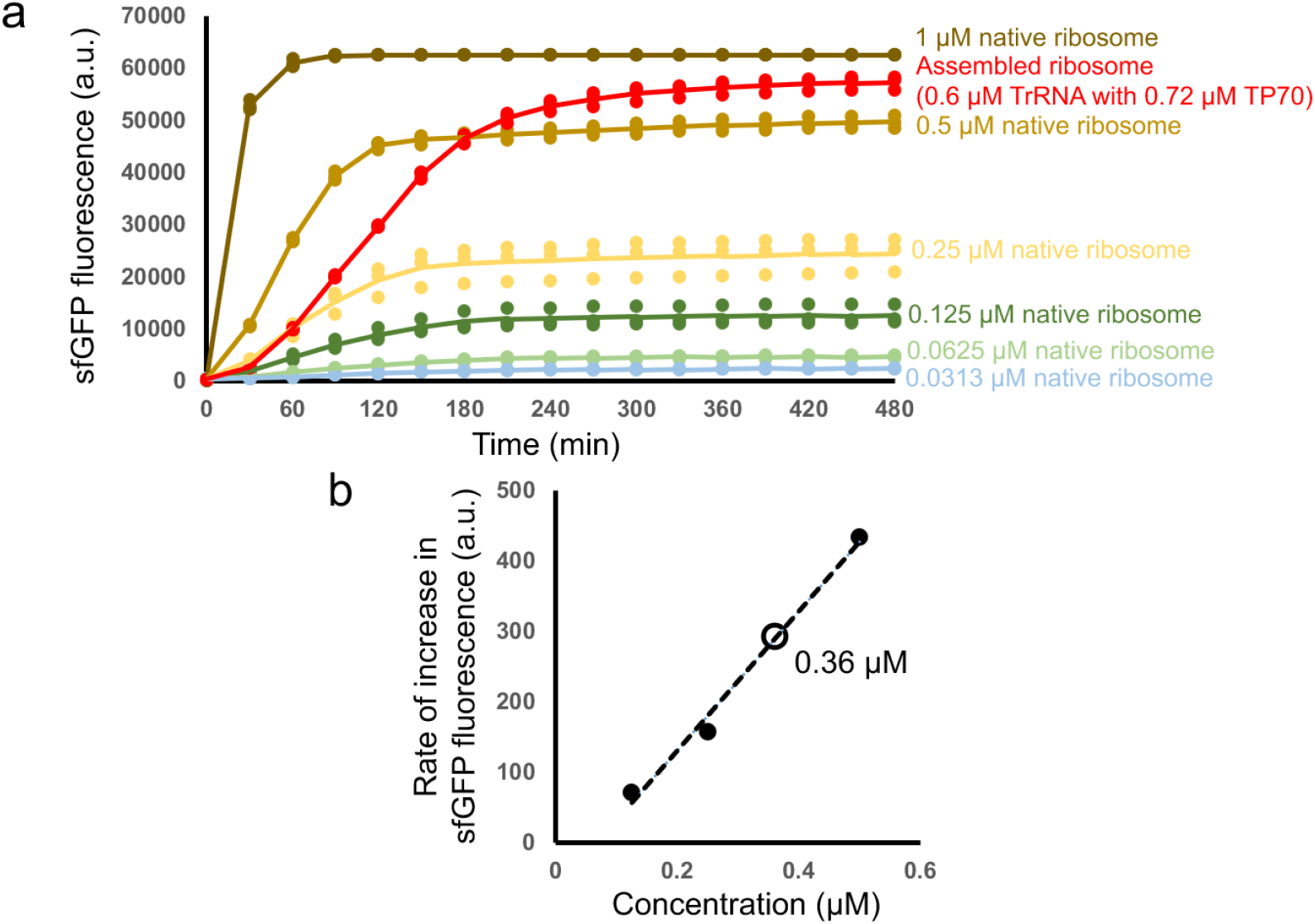
Rate of increase in fluorescence of synthesized sfGFP. (a) sfGFP synthesis was measured in the coupled system with varying concentrations of native ribosome. (b) The estimated concentration of assembled ribosomes was calculated from the native ribosome data. Black dots represent data of the native ribosome and a white dot represent the data of the assembled ribosome.

**Figure 7.**
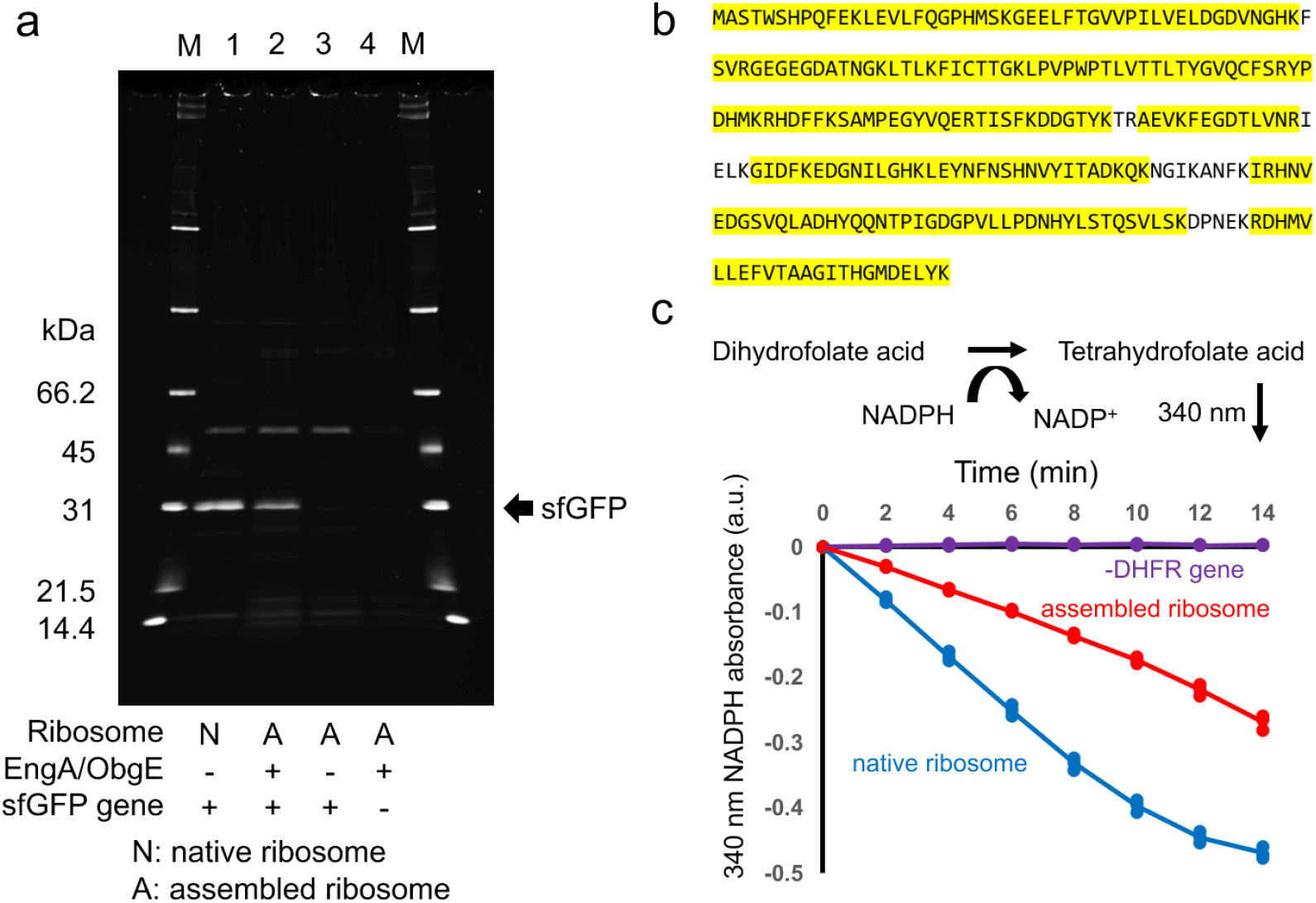
Characterisitics of the synthesized proteins using the assembled ribosome. (a) SDS-PAGE analysis of purified sfGFP. sfGFP synthesis was performed with native ribosome (lane 1) and assembled ribosome (lane 2-4). sfGFP was not synthesized in the absence of EngA and ObgE (lane 3) and in the absence of sfGFP gene (lane 4). M represents the protein size markers. (b) Mass spectrometry analysis of the purified sfGFP. Amino acid sequence of the synthesized sfGFP is shown and the MS-detected sequences are marked with yellow. (c) Activity measurement of the synthesized DHFR. DHFR catalyzes reduction of dihydrofolate acid into tetrahydrofolate acid, in which the reaction could be monitored through reduction of 340 nm NADPH absorbance. Activities of synthesized DHFR with native ribosome and assembled ribosome was measured.

### EngA and ObgE are required for the reassembly of unfolded ribosomes

It has been shown that adding EDTA to ribosomes collapses their structure to a level detectable by electron microscopy^30^. The rRNA used in this study was extracted from native ribosomes with phenol, and it is possible that the partial structure, formed depending on magnesium ion, was still retained, potentially reducing ObgE dependency. We added EDTA to native ribosomes to collapse their structure and then restored the environment by adding magnesium ions. We tested whether ribosome activity would be restored in the presence of ObgE and EngA. The results showed that active ribosomes were formed only in the presence of both factors (**Fig. 8**), suggesting that both ObgE and EngA are equally necessary for reassembling ribosomes.

**Figure 8.**
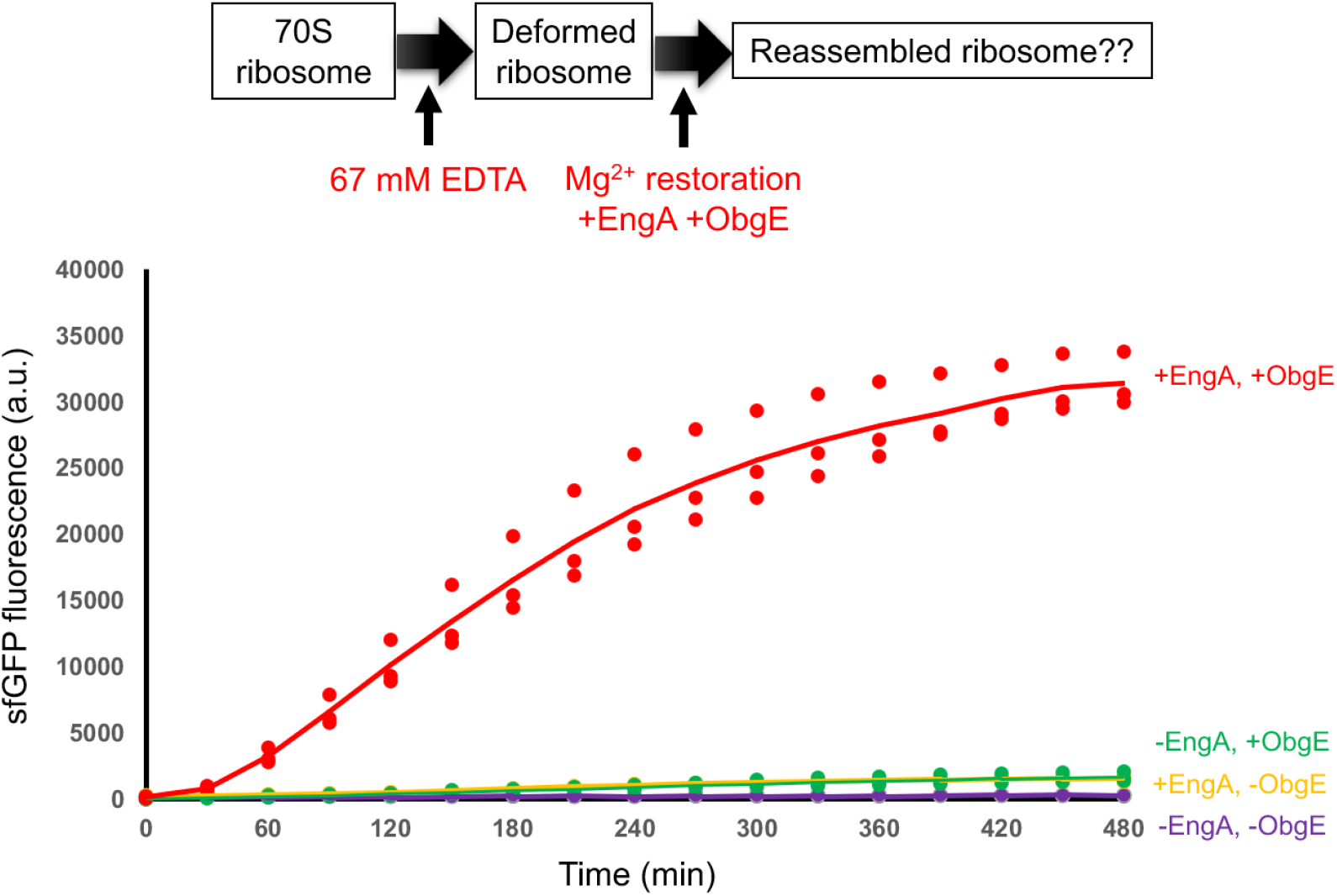
Reassembly of unfolded ribosomes using EngA and ObgE. Ribosomes were unfolded by the addition of high concentration of EDTA and then restored the environment by the addition of magnesium ions. Followingly, reassembly of the ribosome was addressed by using EngA and ObgE. Time course of sfGFP fluorescence are shown in the presence or absence of EngA and ObgE..

## Discussion

In the present study, the change in divalent ion concentration was crucial for the translationally active ribosome assembly (**Fig. 1**). In contrast, in the presence of GTPase factors EngA and ObgE, ribosome assembly and translation reactions could be coupled under constant magnesium ion concentrations (8 mM), indicating the significant role of these two factors under near-physiological conditions (**Fig. 2-7**). Although the dependency on ObgE was slightly mitigated in these experiments, both factors were indispensable for the reversible unfolding and regeneration of ribosomes by manipulation of EDTA and magnesium ion concentrations (**Fig. 8**), suggesting that both of two factors are essential in ribosome biogenesis, compensating for the divalent ion concentration and temperature changes in the conventional in vitro assembly conditions. It is possible that some tertiary structures completed by ObgE catalysis might have been retained in phenol-extracted rRNA prepared from native ribosomes.

The two GTPases identified in this study belong to the 13 highly conserved GTPases in bacteria, implying their significant importance for cell viability^39^. GTPases EngB, HflX, and LepA, examined in this study, also belong to the 13 members and have been suggested to be involved in ribosome assembly^33-35^, whereas no significant activity was detected in the present study. This does not deny their involvement in ribosome assembly under different assembly conditions, and further verification is necessary. We note that an important finding regarding HflX function, which is different from ribosome assembly pathway, has been found very recently^40^. BipA is not included in the 13 members but is reported as a homolog of EF-G and LepA and reported to be involved in the ribosome assembly process^32^, and so it was also tested in this study.

The finding that the two GTPases compensate for the changes in temperature and divalent ion concentration provides new insights into the 50S subunit assembly process. These changes are known to be required for late-stage assembly processes^26^ and therefore, it is suggested that the two factors catalyze the late-stage assembly processes. Indeed, cryo-EM based studies have suggested the involvement of these two factors in the late-stage assembly processes^5^. The structures of *E. coli* 50S subunits with EngA^27^ and ObgE^29^ have been individually elucidated, and structural analysis of 50S subunits in the assembly process obtained by affinity purification with tagged ObgE has been performed^28^. Additionally, in the cryo-EM study on the assembly process of the 50S subunit of *Bacillus subtilis*, a deep relationship with YphC, a homolog of EngA, was demonstrated^33^. These reports suggest the involvement of the two factors in the maturation of critical structures in the late-stage process, including functional core (FC) and peptidyl transferase center (PTC), and the binding of ribosomal proteins involved in the late-stage processes like uL16 and bL36. The present study clarifies the deep relationship between these critical maturation steps in the late-stage processes and the essential operations in the in vitro assembly process identified by Nierhaus and Dohme^25^.

In addition to the structural analyses, genetic studies have suggested the involvement of both proteins in the 50S assembly process^41,42^. Interestingly, both proteins have been reported as suppressor proteins of cold-sensitive RlmE (RrmJ) knockout mutants^43^. RlmE is an enzyme that methylates U2552 in domain V of 23S rRNA, where PTC is located. The 50S subunits obtained from the RlmE-knockout mutants show some translational defects and structural differences, particularly in the FC and PTC regions, compared to the wild-type subunits^44^. Since the cell growth of RlmE knockout mutants restores with increased expression of EngA and ObgE^43^, the structural differences in ribosomes may also be recovered by these factors. This suggests that one of the roles of these two factors includes the maturation of these regions, and their roles might overlap.

However, the present analysis shows that active ribosomes are not assembled with ObgE alone, and the activity of ribosomes assembled with EngA alone is significantly lower than when both factors are used (**Fig. 5**). Furthermore, the regeneration experiment of ribosome structure by EDTA chelation of magnesium ions and subsequent restoration with magnesium ions revealed that the presence of both factors is essential (**Fig. 8**). This suggests that these two factors individually play different roles in the late-stage assembly process. There has been no structural study focusing on both EngA and ObgE at the same time. According to the individual cryo-EM studies, a C-terminal GTPase domain of ObgE binds to the GTPase-associated center (GAC), and its N-terminal domain binds to the interface region towards the PTC from the A site space^28,29^. In contrast, an N-terminal GTPase domain (GD1) of EngA binds near the L1 stalk, its C-terminal GTPase domain (GD2) binds to the P site space, and its C-terminal KH domain binds to the interface region towards the PTC^27^. The N-terminal domain of ObgE and the C-terminal region of EngA, which is not fully observed with electron density maps, are very close, suggesting that they might not bind to the subunit simultaneously. Moreover, the complex obtained by affinity purification with tagged ObgE shows additional electron density maps corresponding to YjgA and RluD, which may prevent EngA binding^28^. Therefore, ObgE and EngA might independently support the late-stage assembly processes in different steps, suggesting that at least two critical maturation steps in the late-stage assembly processes exist. Further structural analysis will elucidate the details of these processes.

Unlike gram-negative bacteria, represented by *E. coli*, gram-positive bacteria, represented by *B. subtilis*, might require a different set of GTPases. Previous reports showed that intermediate 45S or 44.5S subunits accumulate in cells lacking YphC (a homolog of EngA), RbgA, and YsxC (a homolog of EngB)^33,45^. These intermediates lack ribosomal proteins binding in the late-stage processes, such as uL16 and bL36, suggesting stalling before critical maturation steps. Although *B. subtilis* contains a gene annotated as *obgE*, this protein does not appear in a series of studies, while RbgA, which has no corresponding protein in *E. coli*, and YsxC, a homolog of EngB, are essential. This suggests that diversity in support systems for critical maturation steps has arisen during evolution.

The cryo-EM structure of 50S subunits during the assembly process, which are affinity-purified with tagged ObgE, suggested the presence of three factors that may work cooperatively with ObgE, including RsfS, YjgA, and RluD^28^. By contrast, the assembly process progresses well only with ObgE and EngA in the present study. Several scenarios can explain these differences. First, it can be considered that the factors identified in the cryo-EM analysis might be supportive but not essential, consistent with the fact that the genes for these factors are not essential for the cell viability. As another possibility, their involvement in rRNA folding can be considered. In the present study, we use rRNA extracted from native ribosomes (**Fig. 1-7**) or unfolded ribosomes with EDTA (**Fig. 8**). The requirement of ObgE is slightly different between these experiments, suggesting different degrees of rRNA folding in both preparations, which might dependent on the presence of magnesium ions. Moreover, it can be considered that secondary structures formed by simple complementary duplex formation are still preserved in both preparations. In contrast, newly transcribed rRNA is produced without any secondary and tertiary structures. We have attempted the 50S subunit assembly with transcribed rRNAs but failed to achieve active subunits with the two factors (data not shown), suggesting further multifaceted mechanisms might be required for the formation of the 50S subunits. They might include rRNA modifications and associated rRNA folding, and the protein group identified by the cryo-EM study may function in such processes. Further in-depth analysis based on the developed assay system will clarify the overall assembly process of the 50S subunits.

The present study reveals that the process corresponding to the changes in temperature and magnesium concentration, identified by Nierhaus and Dohme^25^, is catalyzed by two GTPases. In reactions using cell extracts like iSAT^11^, the presence of these two factors in the extracts may have facilitated the ribosome assembly without any changes in temperature and ion concentration. While improvement of the iSAT is extensively performed^46-48^, the addition of two factors identified in this study may contribute to such studies. The two GTPases may also be one of the key factors for the coupling systems using cell-free systems for expressing ribosomal proteins^18,20^, as well as the future developments towards self-replicating synthetic cells^9,10^. At the same time, investigation of the mechanisms of actions of the two GTPases, through structural analysis using cryo-EM, protein composition analysis using qMS, and dynamic analysis using imaging techniques will lead to a deeper understanding of ribosome assembly mechanisms.

It should be emphasized that the two factors presented here are not necessary and sufficient factors for the total ribosome assembly. To achieve true bottom-up construction of life represented by creating self-replicating synthetic cells^9,10^, it will be necessary to move beyond components derived from native ribosomes, as used in this study, and instead, employ individually prepared synthetic parts. For example, it will be important to integrate approaches such as ribosome reconstitution from transcribed 23S rRNA^22,23^ and reconstitution using synthetic ribosomal proteins^14-17^, which represents a key future perspective of this study. To accomplish this, continuing efforts using the PURE system-based approaches may help identify additional essential factors, such as rRNA modification enzymes and other biogenesis factors beyond the two factors described here. Through these processes, we also expect to gain a deeper understanding of ribosome assembly mechanisms.

## Methods

### Sample preparation for in vitro experiments

Native ribosomes and the PURE system components were prepared as described previously^49^. Preparation of TP70 was performed according to the conventional methods^31^, as follows: First, 3.5 mL of Buffer 3 (20 mM Hepes-KOH, pH 7.6, 4 mM MgCl_2_, 30 mM NH_4_Cl, 2 mM spermidine, 0.2 mM spermine, 7 mM 2-mercaptoethanol) was added to 1 mL of 24 µM ribosome solution, followed by the addition of 450 µL (0.1 volume) of 1 M Mg(OAc)_2_. Then, 10 mL of glacial acetic acid was added and the resultant solution was stirred at 4°C for 45 minutes. The solution was centrifugated at 10,000 *g* for 30 minutes and the supernatant was recovered. Five volumes of ice-cold acetone (e.g., 75 mL to ∼14.95 mL of solution) was added and the solution was kept at -30°C overnight. The solution was centrifuged at 10,000 *g* for 30 minutes, and then the pellet was vacuum-dried and dissolved in 2 mL of Buffer 5 (20 mM Tris-HCl, pH 7.6, 4 mM Mg(OAc)_2_, 400 mM NH_4_Cl, 6 M urea, 0.2 mM EDTA, 7 mM 2-mercaptoethanol). The solution was dialyzed overnight against 500 mL of Buffer 5 using 3,500 MWCO dialysis membrane. It was further dialyzed against 500 mL of Buffer 4 (20 mM Tris-HCl, pH 7.6, 4 mM Mg(OAc)_2_, 400 mM NH_4_Cl, 0.2 mM EDTA, 7 mM 2-mercaptoethanol) for 45 minutes, three times, using the same membrane. Finally, the concentration was measured and stored in small aliquots at -80°C. Total ribosomal RNAs containing 5S, 16S, and 23S rRNAs Ribosomal RNAs were prepared from the native ribosomes via water-saturated phenol extraction followed by 2-propanol precipitation, according to the same literature^31^. Ribosome biogenesis factors used in this study were prepared as follows. *E. coli* genes for BipA, EngB, HflX, and LepA were cloned into pET15b (Merck Millipore, USA) as His-tag fusion protein. The genes for EngA and ObgE were cloned into pET15b (Merck Millipore, USA) as small ubiquitin-like modifier (SUMO) protein-fusion proteins where His-tag, SUMO protein, and EngA or ObgE were tandemly arranged. Detailed sequences are described in the supplementary data (**Supplementary Data 1**). BipA, EngB, HflX, and LepA proteins were purified in a same way as His-tagged translation factors^49^. EngA and ObgE were purified in a same way as uS2 according to our previous study^15^. DNA templates for the sfGFP synthesis were prepared as previously described^16^.

### Ribosome assembly with a two-step procedure

Reaction mixtures (25 µL) contained 20 mM Tris-HCl, pH 7.4, 4 mM Mg(OAc)_2_, 400 mM NH_4_Cl, 7 mM 2-mercaptoethanol, 22.5 pmol total rRNAs, and 27 pmol TP70. Reactions were carried out at 44 °C for 20 min and then 1 µL of 400 mM Mg(OAc)_2_ solution or water was added to change the magnesium ion concentration. The mixtures were further incubated at 44 °C or 50 °C for 90 min. The mixtures were centrifugated at 5,000 *g* for 5 min at 25 °C and then supernatants were recovered. The assembled ribosomes were concentrated and buffer exchanged using 0.5 mL centrifugal filter units MWCO 30 kDa with ribosome buffer (10 mM Hepes-KOH, pH 7.6, 10 mM Mg(OAc)_2_, 30 mM K(OAc), and 1 mM DTT). The activity of ribosomes was monitored in the home-made PURE system^49^, where reaction mixtures (20 µL) containing the assembled ribosomes from 25 µL mixtures and 5 nM DNA template encoding sfGFP were incubated at 37 °C in Mx3005P (Agilent Technologies, USA), and sfGFP fluorescence was monitored during incubation.

### Ribosome assembly with ribosome biogenesis factors

The assembly reactions were performed based on the PURE system reaction mixtures^49^, in anticipation of future integration with a cell-free translation system. The reaction mixtures (25 µL) contained 50 mM Hepes-KOH, pH7.6, 2 mM spermidine, 1 mM DTT, 2 mM ATP, 2 mM GTP, 1 mM UTP, 1 mM CTP, 10 mM creatine phosphate, 0.1 mM each 20 amino acids, 0.25 µg each of creatine kinase, nucleotide diphosphate kinase, BipA, EngA, EngB, HflX, LepA, and ObgE, varying concentrations of Mg(OAc)_2_ and potassium glutamate, 22.5 pmol total rRNAs, and 27 pmol TP70. The mixtures were incubated at 37 °C for 110 min and they were centrifugated at 5,000 *g* for 5 min at 25 °C, followed by the recovery of supernatants. The assembled ribosomes were concentrated and buffer exchanged using 0.5 mL centrifugal filter units MWCO 30 kDa with ribosome buffer. The activity of ribosomes was monitored in the home-made PURE system^49^, where reaction mixtures (20 µL) containing the assembled ribosomes from 25 µL mixtures and 5 nM DNA template encoding sfGFP were incubated at 37 °C in Mx3005P (Agilent Technologies, USA), and sfGFP fluorescence was monitored during incubation.

### Integration of ribosome assembly and translation systems

Reaction mixtures (20 µL) were based on the home-made PURE system^49^ without ribosomes. In addition to the standard components, 0.25 µg each of BipA, EngA, EngB, HflX, LepA, and ObgE, 22.5 pmol total rRNAs, and 27 pmol TP70 were added in to the mixtures. Concentrations of Mg(OAc)_2_ and potassium glutamate are varied according to the experiments and designated in the figure legends. After the addition of 5 nM DNA template encoding sfGFP, the mixtures were incubated at 37 °C in Mx3005P (Agilent Technologies, USA), and sfGFP fluorescence was monitored during incubation.

### Characterization of the translation products

To compare the protein synthesis efficiency of reconstituted ribosome, 10 μl reactions with varying concentrations of native ribosome (1 μM, 0.5 μM, 0.25 μM, 0.125 μM, 0.0625 μM and 0.03125 μM) and 10 μl ribosome reconstitution reaction (0.72 μM of TP70 with 0.6 μM total rRNAs) were prepared. Reactions were carried out at 37 °C for in Mx3005P (Agilent Technologies, USA) and sfGFP fluorescence was monitored during incubation. For the SDS-PAGE analysis and mass spectrometry analysis of the synthesized sfGFP, Strep-tag II was fused at the N-terminus of sfGFP for the purification^50^. Tagged sfGFP was purified using MagStrep-type 2HC beads (IBA Lifesciences, Germany) according to the manufacturer’s protocol. Elution was performed with the solution containing 2.5 mM D-desthiobiotin and analyzed. SDS-PAGE gel was stained with SYPRO Ruby Protein Gel Stain (Thermo scientfific, USA) according to the manufacturer’s instructions. The activity of synthesized DHFR was measured as follows: DHFR assay reaction mixtures (1 mL) contained 50 mM MES-KOH, pH 7.0, 25 mM Tris-HCl, pH 7.0, 25 mM ethanolamine, 100 mM NaCl, 10 mM 2-mercaptoethanol, 0.1 mM EDTA, 100 μM dihydrofolic acid, and a 5 μL aliquot of the PURE reaction mixture, and they were incubated at 37°C for 15 min. Next, 20 mM NADPH was added to a final concentration of 200 μM. Reaction was aliquoted to a volume of 300 µl on the 96-well titer plate and the decrease in absorbance at 340 nm was measured at 37 °C with Multiskan Go (Thermo scientific, USA).

### Mass spectrometry analysis

The proteins were reduced with 10 mM TCEP at 37 °C for 30 min, alkylated with 20 mM iodoacetamide at 37 °C for 30 min, and quenched with 20 mM L-cysteine. Protein digestion was performed by adding 100 ng of Trypsin/Lys-C mix (Promega, USA) and incubating overnight at 37 °C. Detergents were precipitated by adding a final concentration of 0.5% TFA, after which the mixture was centrifuged at 15,000 *g* for 5 min. The resulting supernatant was purified using GL-Tip SDB (GL Sciences) according to the manufacturer’s instructions and then dried under reduced pressure. The dried peptides were dissolved in 20 μL of 0.1% TFA, and 4 μL of the sample was analyzed by LC-MS. The peptides were concentrated and separated using a nano-LC system (UltiMate 3000, Thermo Scientific, Germany) equipped with a trap column (C18, 0.075 × 20 mm, 3 µm, Acclaim PepMap 100, Thermo Scientific) and a nanocapillary analytical column (C18, 0.075 × 150 mm, 3 µm, Nikkyo Technos, Tokyo, Japan) at a flow rate of 200 nL/min. Mobile phases A (0.1% formic acid) and B (acetonitrile and 0.1% formic acid) were combined in a gradient as follows: 5% B for 5 min, 5–40% B for 75 min, 40–90% B for 1 min, and 90% B for 4 min. MS analysis was performed using an Orbitrap mass spectrometer (Q Exactive, Thermo Scientific) equipped with a nanospray ion source (Nanospray Flex, Thermo Scientific) with the following parameters: spray voltage of 2.2 kV, positive mode, scan range of m/z 310–2000, and 70,000 resolution. The ten most intense multiply charged ions (z = 2–4) were fragmented in the collision cell by collision-induced dissociation (CID). Raw data was processed using Proteome Discoverer 3.2 (Thermo Scientific).

### Reassembly of unfolded ribosomes

First, 0.5 µL of 5 µM ribosome (0.5 pmol) was mixed with 1 µL of 100 mM EDTA (final concentration was 67 mM) and incubated on ice for 30 min. Then, PURE translation mixtures were prepared. In this experiment, we used solution II (Enzyme Mix) of PUREfrex 2.0 (GeneFrontier Corporation) instead of using home-made translation factors and enzymes, whereas other components were based on the home-made PURE system^49^. In addition to the standard components, the mixtures (10 µL) contained 1.5 µL ribosome solution (0.5 pmol in 67 mM EDTA), 0.5 µL of solution II of PUREfrex 2.0, 0.2 µg each of EngA and ObgE. The concentration of potassium glutamate was 150 mM and the concentration of Mg(OAc)_2_ was 32 mM to restore the actual magnesium ion concentration. After the addition of 5 nM DNA template encoding sfGFP, the mixtures were incubated at 37 °C in Mx3005P (Agilent Technologies, USA), and sfGFP fluorescence was monitored during incubation.

## Acknowledgments

This work was supported by CREST (JPMJCR20S4 to YShimizu) from Japan Science and Technology Agency (JST), a Grant-in-Aid (20H05701 and 22H05152 to YShimizu) from the Japan Society for the Promotion of Science (JSPS), the Human Frontier Science Program (RGP0043/2017 to YShimizu), and an intramural Grant-in-Aid from the RIKEN Center for Biosystems Dynamics Research (to YShimizu).

## Author Contributions

Y.Shimizu. designed the study. A.S., W.Y.L., Y.Sakai. and Y.Shimizu. wrote the manuscript. A.S., W.Y.L., Y.Sakai, and K.M. performed the experiments. All authors discussed the results and commented on the manuscript.

