## Supplementary Data 1 for "Single-step in vitro ribosome reconstitution mediated by two GTPase factors, EngA and ObgE"

### Supplementary Data 1. DNA sequences of the plasmids used in this study.

Underlined: T7 promoter or T7 terminator sequence.
Blue: leader sequence containing His-tag.
Green: SUMO protein.
Red: ribosome biogenesis factors.
Purple: ORF of tagged sfGFP and DHFR

>BipA with His-tag in pET15b
TAATACGACTCACTATAGGGGAATTGTGAGCGGATAACAATTCCCCTCTAGAAATAATTTTGTTTAACTTTAA
GAAGGAGATATACCATGGGCAGCAGCCATCATCATCATCATCACAGCAGCGGCCTGGTGCCGCGCGGCAGCCA
TATGATCGAAAAATTGCGTAATATCGCCATCATCGCGCACGTAGACCATGGTAAAACCACCCTGGTAGACAAG
CTGCTCCAACAATCCGGTACGTTCGACTCTCGTGCCGAAACCCAAGAGCGCGTGATGGACTCCAACGATTTGG
AGAAAGAGCGTGGGATTACCATCCTCGCGAAAAACACCGCTATCAAATGGAATGATTACCGTATCAACATCGT
TGATACCCCGGGGCACGCCGACTTCGGTGGTGAAGTTGAACGTGTAATGTCCATGGTAGACTCAGTGCTGCTG
GTGGTTGACGCATTTGACGGCCCGATGCCGCAAACGCGCTTCGTAACCAAAAAAGCGTTTGCTTACGGCCTGA
AGCCGATTGTTGTTATCAACAAAGTTGACCGCCCTGGCGCGCGTCCTGATTGGGTTGTGGATCAGGTATTCGA
TCTGTTCGTTAACCTCGACGCGACCGACGAGCAGCTGGACTTCCCGATCGTTTACGCTTCTGCGCTGAACGGT
ATCGCGGGTCTGGACCACGAAGATATGGCGGAAGACATGACCCCGCTGTACCAGGCGATTGTTGACCACGTTC
CTGCGCCGGACGTTGACCTTGACGGTCCGTTCCAGATGCAGATTTCTCAGCTCGATTACAACAGCTATGTTGG
CGTTATCGGCATTGGCCGCATCAAGCGCGGTAAAGTGAAGCCGAACCAGCAGGTCACTATCATCGATAGCGAA
GGCAAAACCCGCAACGCGAAAGTCGGTAAAGTGCTGGGCCACCTCGGTCTGGAACGTATCGAAACCGATCTGG
CGGAAGCTGGCGATATCGTTGCGATCACGGGCCTTGGCGAACTGAACATTTCTGACACCGTTTGCGACACGCA
AAACGTTGAAGCGCTGCCGGCACTCTCCGTTGATGAGCCGACCGTTTCTATGTTCTTCTGCGTTAACACCTCG
CCGTTCTGCGGTAAAGAAGGTAAGTTCGTAACGTCTCGTCAGATCCTGGATCGTCTGAACAAAGAACTGGTAC
ACAACGTTGCGCTGCGCGTAGAAGAAACCGAAGACGCCGATGCGTTCCGCGTTTCTGGTCGTGGCGAACTGCA
CCTGTCTGTTCTGATCGAAAACATGCGTCGTGAAGGTTTCGAACTGGCGGTATCCCGTCCGAAAGTTATCTTC
CGTGAAATCGACGGTCGTAAACAAGAGCCGTATGAAAACGTGACTCTGGACGTTGAAGAACAGCATCAGGGTT
CTGTAATGCAGGCGCTGGGCGAACGTAAAGGCGACCTGAAAAACATGAATCCAGACGGTAAAGGCCGCGTACG
TCTCGACTACGTGATCCCAAGCCGTGGTCTGATTGGCTTCCGTTCTGAGTTCATGACCATGACTTCCGGTACT
GGTCTGCTGTACTCCACCTTCAGCCACTACGACGACGTACGTCCGGGTGAAGTGGGTCAGCGTCAGAACGGCG
TACTGATCTCTAACGGTCAGGGTAAAGCGGTCGCGTTCGCGCTGTTCGGTCTGCAGGATCGCGGTAAGCTGTT
CCTCGGTCACGGTGCAGAAGTTTACGAAGGTCAGATTATCGGTATTCATAGCCGCTCTAACGACCTGACTGTA
AACTGCCTGACCGGTAAGAAACTGACCAACATGCGTGCTTCCGGTACTGACGAAGCCGTTGTTCTGGTTCCGC
CTATCCGCATGACTCTGGAACAAGCTCTGGAGTTCATCGATGATGACGAACTGGTAGAAGTGACTCCGACCTC
TATCCGTATTCGTAAACGTCACCTGACGGAAAACGATCGTCGCCGCGCCAACCGCGCACCGAAAGACGATTAA
GGATCCGGCTGCTAACAAAGCCCGAAAGGAAGCTGAGTTGGCTGCTGCCACCGCTGAGCAATAACTAGCATAA
CCCCTTGGGGCCTCTAAACGGGTCTTGAGGGGTTTTTTG

>EngB with His-tag in pET15b
TAATACGACTCACTATAGGGGAATTGTGAGCGGATAACAATTCCCCTCTAGAAATAATTTTGTTTAACTTTAA
GAAGGAGATATACCATGGGCAGCAGCCATCATCATCATCATCACAGCAGCGGCCTGGTGCCGCGCGGCAGCCA

TATGACTAATTTGAATTATCAACAGACGCATTTTGTGATGAGTGCGCCTGATATTCGCCACCTACCTTCCGAT
ACCGGAATTGAAGTGGCTTTTGCAGGCCGTTCCAACGCAGGTAAATCCAGCGCGCTGAACACGCTGACTAACC
AGAAAAGCCTGGCTCGTACCTCAAAAACCCCAGGGCGCACCCAGCTTATCAACCTGTTTGAAGTGGCTGACGG
CAAGCGTCTGGTTGACTTGCCTGGGTACGGTTATGCGGAAGTCCCGGAAGAGATGAAGCGCAAATGGCAGCGT
GCGCTCGGCGAATACCTCGAAAAACGTCAGAGCCTGCAAGGTCTGGTGGTGCTAATGGATATTCGCCATCCGC
TGAAAGATTTGGATCAGCAGATGATTGAGTGGGCGGTAGACAGCAATATCGCCGTTCTGGTGCTGCTGACCAA
AGCGGACAAACTGGCAAGCGGCGCACGTAAAGCGCAATTGAATATGGTGCGTGAAGCTGTACTGGCGTTTAAC
GGTGATGTGCAGGTTGAAACGTTTTCTTCGTTGAAGAAACAAGGCGTGGACAAGCTGCGGCAGAAACTGGATA
CCTGGTTTAGCGAGATGCAGCCTGTAGAAGAAACGCAGGACGGCGAATAAGGATCCGGCTGCTAACAAAGCCC
GAAAGGAAGCTGAGTTGGCTGCTGCCACCGCTGAGCAATAACTAGCATAACCCCTTGGGGCCTCTAAACGGGT
CTTGAGGGGTTTTTTG

>HflX with His-tag in pET15b
TAATACGACTCACTATAGGGGAATTGTGAGCGGATAACAATTCCCCTCTAGAAATAATTTTGTTTAACTTTAA
GAAGGAGATATACCATGGGCAGCAGCCATCATCATCATCATCACAGCAGCGGCCTGGTGCCGCGCGGCAGCCA
TATGTTTGACCGTTATGATGCTGGTGAGCAGGCGGTACTGGTACACATCTATTTTACGCAAGACAAAGATATG
GAAGACCTCCAGGAGTTTGAATCTCTGGTCTCTTCCGCCGGTGTCGAAGCATTGCAGGTGATTACCGGTAGCC
GTAAAGCGCCGCACCCAAAGTATTTTGTAGGTGAAGGTAAAGCAGTTGAAATTGCGGAAGCTGTCAAAGCGAC
GGGTGCTTCGGTCGTTCTTTTTGACCATGCCCTGAGCCCGGCGCAAGAGCGTAACCTGGAGCGTTTGTGCGAG
TGTCGTGTTATCGACCGCACCGGCCTTATTTTAGATATTTTCGCCCAACGTGCGCGTACCCATGAGGGTAAGT
TGCAGGTTGAGCTGGCGCAGCTGCGCCATCTGGCTACGCGCCTGGTGCGTGGCTGGACCCACCTTGAAAGACA
GAAAGGCGGGATAGGTTTGCGTGGTCCGGGTGAAACCCAGCTCGAAACCGACCGTCGTTTGTTGCGTAATCGC
ATCGTGCAGATACAGTCGCGCCTGGAAAGAGTTGAAAAGCAGCGTGAGCAGGGGCGGCAATCGCGTATCAAAG
CCGACGTTCCTACTGTTTCGCTGGTGGGATATACCAACGCCGGTAAATCTACCCTTTTCAATCGCATCACCGA
AGCGCGGGTCTACGCGGCAGACCAGTTGTTTGCCACCCTCGACCCGACGTTGCGGCGTATTGACGTTGCAGAT
GTCGGTGAAACCGTACTTGCAGATACCGTAGGGTTTATTCGCCACCTGCCGCACGATCTGGTGGCGGCATTTA
AAGCCACGTTACAAGAGACGCGGCAAGCCACATTACTGCTGCACGTCATTGATGCGGCGGATGTGCGTGTACA
AGAAAACATCGAAGCGGTGAATACGGTTCTTGAAGAGATCGACGCTCACGAGATCCCAACCCTGCTGGTGATG
AACAAGATCGATATGCTGGAAGATTTCGAACCGCGTATTGATCGGGACGAAGAGAACAAACCGAACCGTGTCT
GGCTTTCCGCACAGACCGGAGCGGGGATACCACAGCTTTTTCAGGCTTTGACGGAGCGGCTTTCCGGCGAGGT
GGCGCAGCATACATTGCGTCTGCCACCGCAGGAAGGGCGTCTGAGAAGTCGTTTTTATCAGCTTCAGGCAATA
GAAAAAGAGTGGATGGAGGAGGACGGCAGCGTAAGTCTGCAAGTTCGTATGCCGATCGTTGACTGGCGTCGCC
TCTGTAAACAAGAACCGGCGTTGATCGATTACCTGATCTAAGGATCCGGCTGCTAACAAAGCCCGAAAGGAAG
CTGAGTTGGCTGCTGCCACCGCTGAGCAATAACTAGCATAACCCCTTGGGGCCTCTAAACGGGTCTTGAGGGG
TTTTTTG

>LepA with His-tag in pET15b
TAATACGACTCACTATAGGGGAATTGTGAGCGGATAACAATTCCCCTCTAGAAATAATTTTGTTTAACTTTAA
GAAGGAGATATACCATGGGCAGCAGCCATCATCATCATCATCACAGCAGCGGCCTGGTGCCGCGCGGCAGCCA
TATGAAGAATATACGTAACTTTTCGATCATAGCTCACATTGACCACGGTAAATCGACGCTGTCTGACCGTATT
ATCCAGATCTGCGGTGGCCTGTCTGACCGTGAAATGGAGGCGCAGGTTCTCGATTCCATGGATCTTGAGCGTG
AGCGTGGCATTACCATCAAAGCGCAAAGCGTGACGCTGGACTACAAAGCGTCTGACGGCGAAACCTATCAGCT
TAACTTTATCGACACCCCGGGCCACGTAGACTTCTCCTATGAAGTTTCCCGTTCGCTGGCTGCCTGTGAAGGT

GCATTGCTGGTGGTCGACGCCGGGCAGGGCGTAGAAGCGCAAACCCTGGCAAACTGCTACACCGCCATGGAAA
TGGATCTCGAAGTTGTGCCGGTACTGAACAAGATTGACCTGCCGGCAGCCGATCCTGAACGCGTGGCGGAAGA
AATTGAAGATATCGTCGGCATCGACGCCACCGACGCGGTGCGCTGTTCAGCGAAAACCGGCGTTGGTGTGCAG
GACGTTCTCGAACGTCTGGTGCGCGACATTCCGCCGCCGGAAGGCGATCCGGAAGGCCCGTTGCAGGCACTAA
TTATCGACTCATGGTTCGACAACTACCTGGGCGTTGTTTCACTTATCCGTATTAAAAACGGCACCCTGCGTAA
GGGCGACAAAGTGAAAGTCATGAGTACCGGGCAGACCTATAACGCCGACCGTCTGGGCATCTTCACGCCGAAA
CAGGTTGACCGCACTGAACTGAAATGTGGCGAAGTAGGCTGGCTCGTATGTGCGATTAAAGATATCCACGGCG
CTCCAGTCGGCGATACCTTAACGCTGGCGCGTAATCCGGCAGAAAAGGCGCTGCCTGGCTTTAAGAAAGTCAA
ACCGCAGGTATACGCCGGTCTGTTCCCGGTAAGTTCCGACGACTATGAAGCCTTCCGTGACGCGCTGGGTAAA
CTCAGCCTGAACGATGCCTCACTGTTCTATGAGCCGGAAAGCTCCAGCGCGCTGGGCTTTGGTTTCCGCTGCG
GCTTCCTCGGCCTGCTGCACATGGAGATCATCCAGGAACGTCTGGAACGTGAATACGATCTGGATCTGATCAC
CACTGCGCCGACCGTAGTGTATGAAGTTGAAACCACGTCAAGAGAAGTTATCTACGTCGACAGCCCATCCAAG
CTGCCTGCGGTAAATAACATCTACGAACTGCGCGAGCCGATTGCAGAGTGTCACATGCTGCTGCCGCAGGCAT
ATCTCGGCAACGTTATTACGTTGTGCGTAGAAAAACGCGGCGTGCAGACCAATATGGTTTACCACGGTAATCA
GGTGGCGCTGACGTACGAGATCCCGATGGCGGAAGTGGTGCTCGACTTCTTCGATCGCCTGAAATCTACCTCG
CGTGGTTATGCGTCTCTGGATTACAACTTCAAGCGCTTCCAGGCGTCCGACATGGTACGTGTAGACGTATTAA
TCAACGGTGAACGTGTTGATGCGCTGGCGTTGATCACCCACCGTGATAATTCGCAAAACCGCGGTCGCGAGTT
GGTGGAGAAGATGAAAGATCTGATCCCACGCCAGCAGTTTGATATCGCCATTCAGGCAGCGATTGGTACGCAC
ATCATTGCGCGATCCACCGTGAAACAGCTGCGTAAAAACGTACTGGCTAAATGTTATGGCGGCGATATCAGCC
GTAAGAAAAAGCTGCTGCAGAAGCAGAAAGAAGGTAAGAAACGCATGAAGCAGATCGGTAACGTCGAGCTGCC
GCAGGAAGCGTTCCTCGCCATTCTGCACGTCGGCAAAGACAACAAATAAGGATCCGGCTGCTAACAAAGCCCG
AAAGGAAGCTGAGTTGGCTGCTGCCACCGCTGAGCAATAACTAGCATAACCCCTTGGGGCCTCTAAACGGGTC
TTGAGGGGTTTTTTG

>EngA fused with His-tagged SUMO protein in pET-15b
TAATACGACTCACTATAGGGGAATTGTGAGCGGATAACAATTCCCCTCTAGAAATAATTTTGTTTAACTTTAA
GAAGGAGATATACCATGGGCAGCAGCCATCATCATCATCATCACAGCAGCGGCATGTCGGACTCAGAAGTCAA
TCAAGAAGCTAAGCCAGAGGTCAAGCCAGAAGTCAAGCCTGAGACTCACATCAATTTAAAGGTGTCCGATGGA
TCTTCAGAGATCTTCTTCAAGATCAAAAAGACCACTCCTTTAAGAAGGCTGATGGAAGCGTTCGCTAAAAGAC
AGGGTAAGGAAATGGACTCCTTAAGATTCTTGTACGACGGTATTAGAATTCAAGCTGATCAGACCCCTGAAGA
TTTGGACATGGAGGATAACGATATTATTGAGGCTCACAGAGAACAGATTGGTGGTATGGTACCTGTGGTCGCG
CTTGTCGGGCGCCCTAACGTAGGAAAATCCACGTTATTTAACCGTCTAACTCGCACCCGAGATGCGCTGGTTG
CGGATTTCCCGGGTCTGACTCGTGACCGTAAGTACGGTCGTGCGGAAATTGAAGGCCGTGAGTTTATCTGTAT
TGATACCGGCGGGATTGATGGCACAGAAGACGGTGTAGAAACCCGCATGGCGGAACAGTCGCTGCTGGCGATT
GAAGAAGCGGACGTCGTACTGTTTATGGTGGATGCGCGCGCGGGCCTGATGCCGGCAGATGAAGCGATTGCCA
AACATCTGCGCTCCCGTGAAAAACCGACCTTCCTGGTGGCAAACAAAACTGACGGTCTGGATCCCGATCAGGC
AGTGGTTGATTTCTACTCGCTTGGTTTAGGTGAAATCTACCCGATCGCCGCGTCTCACGGTCGTGGCGTATTA
AGTCTGCTGGAGCATGTGCTGCTGCCGTGGATGGAAGATCTCGCACCGCAAGAGGAAGTCGACGAAGACGCTG
AATACTGGGCGCAATTTGAAGCGGAAGAGAACGGCGAAGAAGAAGAGGAAGACGACTTCGACCCGCAAAGTCT
GCCGATCAAACTGGCGATTGTGGGTCGTCCGAACGTAGGTAAGTCTACACTCACTAACCGTATTCTTGGTGAA
GAGCGCGTTGTTGTTTACGACATGCCTGGCACGACGCGTGACAGCATCTACATCCCAATGGAACGCGATGGAC
GTGAGTATGTGCTCATTGACACCGCTGGCGTACGTAAACGCGGCAAAATCACCGATGCTGTAGAGAAATTCTC
CGTAATCAAAACGTTGCAGGCCATTGAAGACGCCAACGTGGTGATGTTAGTGATTGATGCGCGCGAAGGTATT

TCCGATCAGGATCTCTCGCTGCTGGGCTTTATTCTCAATAGTGGGCGCTCACTTGTCATTGTGGTGAATAAGT
GGGATGGCCTGAGTCAGGAAGTGAAAGAGCAGGTGAAAGAAACGCTGGACTTCCGTCTGGGCTTTATCGATTT
TGCTCGTGTGCACTTTATCTCTGCCTTGCACGGCAGTGGTGTTGGTAACTTGTTTGAATCAGTACGTGAAGCG
TATGACAGCTCCACCCGTCGTGTGGGGACCTCTATGCTGACGCGCATCATGACGATGGCTGTTGAAGATCACC
AACCGCCGCTGGTACGCGGTCGTCGTGTGAAGCTGAAATATGCCCACGCCGGTGGTTATAACCCGCCGATTGT
GGTGATTCACGGTAATCAGGTGAAAGACCTGCCTGATTCCTACAAGCGCTACTTGATGAACTACTTCCGCAAA
TCGCTGGACGTAATGGGATCGCCGATTCGTATTCAGTTCAAAGAAGGGGAAAACCCGTATGCGAATAAGCGTA
ACACCCTGACGCCAACCCAGATGCGTAAACGTAAGCGTCTGATGAAGCACATCAAGAAAAATAAATAACATAT
GCCCGGGCTCGAGGGACCCCGCGGGCGGCCGCGTCGACGGATCCGGCTGCTAACAAAGCCCGAAAGGAAGCTG
AGTTGGCTGCTGCCACCGCTGAGCAATAACTAGCATAACCCCTTGGGGCCTCTAAACGGGTCTTGAGGGGTTT
TTTG

>ObgE fused with His-tagged SUMO protein in pET-15b

TAATACGACTCACTATAGGGGAATTGTGAGCGGATAACAATTCCCCTCTAGAAATAATTTTGTTTAACTTTAA
GAAGGAGATATACCATGGGCAGCAGCCATCATCATCATCATCACAGCAGCGGCATGTCGGACTCAGAAGTCAA
TCAAGAAGCTAAGCCAGAGGTCAAGCCAGAAGTCAAGCCTGAGACTCACATCAATTTAAAGGTGTCCGATGGA
TCTTCAGAGATCTTCTTCAAGATCAAAAAGACCACTCCTTTAAGAAGGCTGATGGAAGCGTTCGCTAAAAGAC
AGGGTAAGGAAATGGACTCCTTAAGATTCTTGTACGACGGTATTAGAATTCAAGCTGATCAGACCCCTGAAGA
TTTGGACATGGAGGATAACGATATTATTGAGGCTCACAGAGAACAGATTGGTGGTATGAAGTTTGTTGATGAA
GCATCGATTCTGGTCGTTGCAGGTGATGGCGGTAATGGTTGCGTGAGCTTCCGCCGCGAAAAGTATATTCCGA
AAGGCGGCCCGGATGGCGGCGACGGCGGTGATGGTGGTGACGTATGGATGGAAGCCGACGAGAACCTGAACAC
GCTTATCGATTATCGTTTTGAAAAATCTTTCCGTGCAGAGCGCGGTCAGAATGGCGCAAGCCGCGACTGTACC
GGTAAGCGCGGTAAAGACGTGACGATTAAAGTGCCGGTAGGTACGCGTGTAATCGACCAGGGTACTGGTGAAA
CCATGGGCGATATGACCAAACACGGTCAGCGTCTGCTGGTTGCTAAGGGCGGCTGGCACGGTCTGGGCAATAC
CCGTTTCAAATCGTCCGTTAACCGTACACCGCGGCAGAAAACCAACGGCACGCCGGGCGATAAGCGCGAGCTG
CTGCTGGAGCTGATGCTGCTGGCTGACGTCGGTATGTTGGGGATGCCAAACGCGGGTAAATCGACCTTTATTC
GTGCGGTATCGGCGGCTAAACCGAAAGTGGCGGATTATCCGTTTACCACTCTGGTGCCAAGTCTGGGTGTGGT
ACGAATGGACAACGAAAAGAGCTTCGTTGTTGCCGATATTCCAGGACTGATTGAAGGCGCTGCGGAAGGCGCA
GGTCTGGGCATTCGCTTCCTGAAGCACCTGGAACGTTGCCGCGTCCTGTTGCACCTCATCGATATCGATCCGA
TTGACGGCACCGATCCGGTTGAAAACGCGCGTATTATTATCAGCGAGCTGGAAAAATACAGCCAGGATCTGGC
GACGAAACCGCGTTGGTTAGTGTTCAACAAGATCGATCTGCTGGATAAGGTAGAAGCCGAAGAGAAAGCGAAA
GCGATCGCTGAGGCGCTGGGCTGGGAAGATAAATATTATCTGATCTCTGCGGCGAGTGGACTGGGCGTGAAAG
ATCTCTGCTGGGATGTGATGACCTTTATCATTGAAAACCCGGTCGTGCAGGCTGAAGAAGCGAAACAGCCAGA
GAAAGTCGAATTCATGTGGGATGATTATCATCGCCAGCAGCTTGAAGAGATTGCTGAAGAGGATGATGAAGAC
TGGGATGACGACTGGGACGAAGACGACGAAGAAGGCGTTGAGTTCATTTACAAGCGTTAACATATGCCCGGGC
TCGAGGGACCCCGCGGGCGGCCGCGTCGACGGATCCGGCTGCTAACAAAGCCCGAAAGGAAGCTGAGTTGGCT
GCTGCCACCGCTGAGCAATAACTAGCATAACCCCTTGGGGCCTCTAAACGGGTCTTGAGGGGTTTTTTG

>sfGFP fused with Strep-tag II in the previously constructed vector based on pUC18 (ref. 50 of the manuscript)

TAATACGACTCACTATAGGGAGACCACAACGGTTTCCCTCTAGAAATAATTTTGTTTAACTTTAAGAAGGAGA
TATACCATGGCTAGCACTTGGTCTCACCCGCAGTTTGAGAAGCTGGAAGTTCTGTTCCAGGGGCCCcatATGA
GTAAAGGAGAAGAACTTTTCACTGGAGTTGTCCCAATTCTTGTTGAATTAGATGGTGATGTTAATGGGCACAA

ATTTTCTGTCCGTGGAGAGGGTGAAGGTGATGCAACAAACGGAAAACTTACCCTTAAATTTATTTGCACTACT
GGAAAACTACCTGTTCCATGGCCAACACTTGTCACTACTTTAACTTATGGTGTTCAATGCTTTTCCCGTTATC
CGGATCACATGAAACGGCATGACTTTTTCAAGAGTGCCATGCCCGAAGGTTATGTACAGGAACGCACTATATC
TTTCAAAGATGACGGGACCTACAAGACGCGTGCTGAAGTCAAGTTTGAAGGTGATACCCTTGTTAATCGTATC
GAGTTAAAAGGTATTGATTTTAAAGAAGATGGAAACATTCTCGGACACAAACTCGAGTACAACTTTAACTCAC
ACAATGTATACATCACGGCAGACAAACAAAAGAATGGAATCAAAGCTAACTTCAAAATTCGCCACAACGTTGA
AGATGGATCCGTTCAACTAGCAGACCATTATCAACAAAATACTCCAATTGGCGATGGCCCTGTCCTTTTACCA
GACAACCATTACCTGTCGACACAATCTGTCCTTTCGAAAGATCCCAACGAAAAGCGTGACCACATGGTCCTTC
TTGAGTTTGTAACTGCTGCTGGGATTACACATGGCATGGATGAGCTCTACAAATAAGGATCCCGGGAATTCTC
GAGAAGCTTTAGCATAACCCCTTGGGGCCTCTAAACGGGTCTTGAGGGGTTTTTTG

>DHFR fused with Strep-tag II in the previously constructed vector based on pUC18 (ref. 50 of the manuscript)

TAATACGACTCACTATAGGGAGACCACAACGGTTTCCCTCTAGAAATAATTTTGTTTAACTTTAAGAAGGAGA
TATACCATGGCTAGCACTTGGTCTCACCCGCAGTTTGAGAAGCTGGAAGTTCTGTTCCAGGGGCCCCATATGA
TCAGTCTGATTGCGGCGTTAGCGGTAGATCGCGTTATCGGCATGGAAAACGCCATGCCGTGGAACCTGCCTGC
CGATCTCGCCTGGTTTAAACGCAACACCTTAAATAAACCCGTGATTATGGGCCGCCATACCTGGGAATCAATC
GGTCGTCCGTTGCCAGGACGCAAAAATATTATCCTCAGCAGTCAACCGGGTACGGACGATCGCGTAACGTGGG
TGAAGTCGGTGGATGAAGCCATCGCGGCGTGTGGTGACGTACCAGAAATCATGGTGATTGGCGGCGGTCGCGT
TTATGAACAGTTCTTGCCAAAAGCGCAAAAACTGTATCTGACGCATATCGACGCAGAAGTGGAAGGCGACACC
CATTTCCCGGATTACGAGCCGGATGACTGGGAATCGGTATTCAGCGAATTCCACGATGCTGATGCGCAGAACT
CTCACAGCTATTGCTTTGAGATTCTGGAGCGGCGGTAAGGATCCCGGGAATTCTCGAGAAGCTTTAGCATAAC
CCCTTGGGGCCTCTAAACGGGTCTTGAGGGGTTTTTTG
